# Vector-Mediated Transport Producing Drug-Like Peptides

**DOI:** 10.1101/507434

**Authors:** Kenneth A. Gruber, James A. Cowan, Ada Cowan, Wenbin Qi, Stephen Pearson, Martin J. Ross, Christine Wachnowsky, Fabio Gallazzi, Shaokai Jiang, Steve R. Van Doren, Michael F. Callahan

**Affiliations:** Tensive Controls, Inc., Columbia, MO 65211; John M. Dalton Cardiovascular Research Center and Department of Medical Pharmacology & Physiology, University of Missouri, Columbia, MO 65212; MetalloPharm, LLC, Columbus, OH; Department of Chemistry and Biochemistry, The Ohio State University, Columbus, OH; Department of Chemistry and Peptide Synthesis Core, University of Missouri, Columbia, MO 65211; Department of Chemistry and NMR Core Facility, University of Missouri, Columbia, MO 65211; Department of Biochemistry, University of Missouri, Columbia, MO 65211

## Abstract

Drugs that are structural mimetics of peptides (e.g. small molecules) have been plagued by problems associated with oral availability and transcellular movement. Vector-mediated transport, where a potentially therapeutic drug is covalently linked to another molecule that is a ligand for an active transport or transcytosis system, was developed as an approach for moving a drug across the blood-brain-barrier. We now report a vector approach that produced peptides with oral activity, blood-brain-barrier transport, and extended *in vivo* half-life. Generating these properties requires secondary structure stabilization into a β hairpin, and the addition of a C-terminal dipeptide sequence composed of non-polar residues. Peptides with biological activity incompatible with these derivatizations were covalently linked to a model transport vector, producing a chimera with the therapeutic activity of the peptide and the transport properties of the vector. Our platform technology may be a general approach for the design of drug-like peptides.

## INTRODUCTION

Biologically active peptides are often used as models for the development of drugs that are mimetics of the peptide’s essential sequence for biological activity [the pharmacophore; (Sun, 2013, Lax, 2010, Pichereau, 2013)]. While peptides are potentially attractive as therapeutic agents due to their high specific activity and low toxicity, their lack of oral activity and transcellular movement, as well as short *in vivo* half-lives have limited their clinical utility (C&EN, 2005). Peptide mimetics (“small molecules”) have also been plagued by unforeseen toxicity and/or off-target side-effects that are rarely encountered with peptides (Lax, 2010, Sun, 2013). Numerous reviews of peptide drug development have called for new chemical approaches to make peptides more drug-like (Brogden and Brogden, 2011, Lax, 2010, Khafagy el and Morishita, 2012).

Reports that an occasional biologically active peptide under investigation had blood-brain-barrier (BBB) transport and/or oral activity for unexplained reasons are scattered throughout the medical research literature (Rodriguez et al., 1993, Sefler et al., 1995, Sutton et al., 2008, Hess et al., 2008). The few attempts to explore this mechanism (or mechanisms) of action have generally failed to define any specific structural features governing these activities, which could then be applied to peptides with potentially therapeutic activity to produce drug-like analogs.(Hess et al., 2007).

We now describe a technological approach that successfully converted several peptides with potentially therapeutic activities into drug-like analogs, by converting them into apparent ligands or “vectors” for a carrier-mediated transport system. Vector-mediated transport, attaching a molecule of interest to a ligand for a transport mechanism, has been used to enhance peptide movement across the BBB (Bockenhoff et al., 2014, Rousselle et al., 2000). We first applied this concept to the development of a melanocortin 4 receptor (MC4R) antagonist peptide to treat the condition of cachexia. Success in this effort resulted in additional applications of the technology to anti-cancer and anti-hepatitis C peptide drug development, with positive results in both efforts. This approach potentially serves as a platform technology for the production of peptide pharmaceuticals.

## RESULTS

### Drug-Like Peptide Melanocortin-4 Receptor Antagonist

Literature analysis showed that many peptides with reported BBB transport, oral activity, or hepatic active transport had common structural features (Table 1). These included cyclization and/or a proline residue within or adjacent to the peptide’s pharmacophore and a C-terminus composed of nonpolar amino acid residues or chemical groups. We produced several MC4R antagonist peptides modeled around common features of the peptides in Table 1; initially peptides # 2, 3, 4 and 5, and demonstrated their anti-cachexia activity to intraperitoneal (IP) LPS challenges using parenteral and oral routes of peptide administration. Similar structural requirements for drug-like activity were subsequently found in peptides # 6-12 (Table 1), which are examples of peptides actively transported by a common carrier system (Bertrams and Ziegler, 1991, Ziegler et al., 1988, Frimmer and Ziegler, 1988).

**Table 1:**
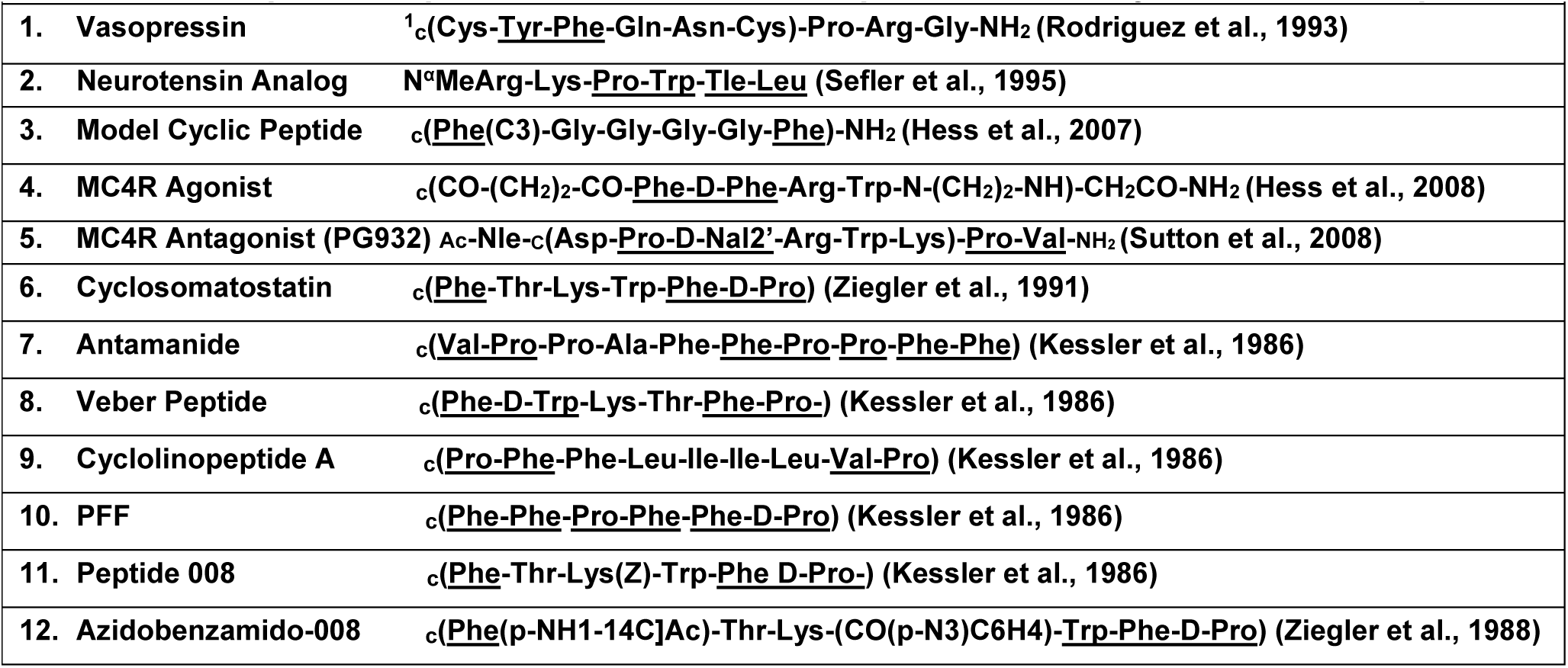
Peptides Reported to Have BBB Transport, Oral Activity or Active Transport. Shows a series of peptide structures reported to have BBB transport, oral activity or active transport (Nal=naphthylalanine). Underlined dipeptide sequences conform to potentially common requirements for transport activity. Essential motifs include at least one Pro-aromatic residue dipeptide and one dipeptide of two nonpolar residues, the latter is ideally D-Val-D-Pro (see TCMCB07 below). As described in Results, the Pro-aromatic motif stabilizes peptides into a β hairpin, while the nonpolar peptide is sequence specific for different forms of transport. Several of the peptides have multiple examples of one or both of these motifs. In six of the 12 peptides, one of the motifs is composed of the notational first and last residues of the depicted cyclized sequence; i.e. adjacent residues in the cyclized peptides. In some instances (peptides #6, 11, 12), the two motifs overlap (e.g. Phe-Pro or Pro-Phe). (See Discussion).

***Ac****-Nle-^1^ c(Asp-Pro-D-Nal2’-Arg-Trp-Lys)-D-Pro-D-Val-****NH*_*2*_**: (TCMCB02): TCMCB02 is an analog of PG932 (Table 1, Peptide #5), with “D” enantiomers of proline and valine substituted in the C-terminal dipeptide to enhance *in vivo* stability (Tugyi et al., 2005), and suppress cardiovascular side-effects (see TCMCB07 below and Discussion). TCMCB02 had significant anti-cachexia activity [measured as increased body weight (BW) and food intake] when given by a parenteral route of administration (Fig. 1A&B), but had no oral anticachexia activity.

**Fig 1.**
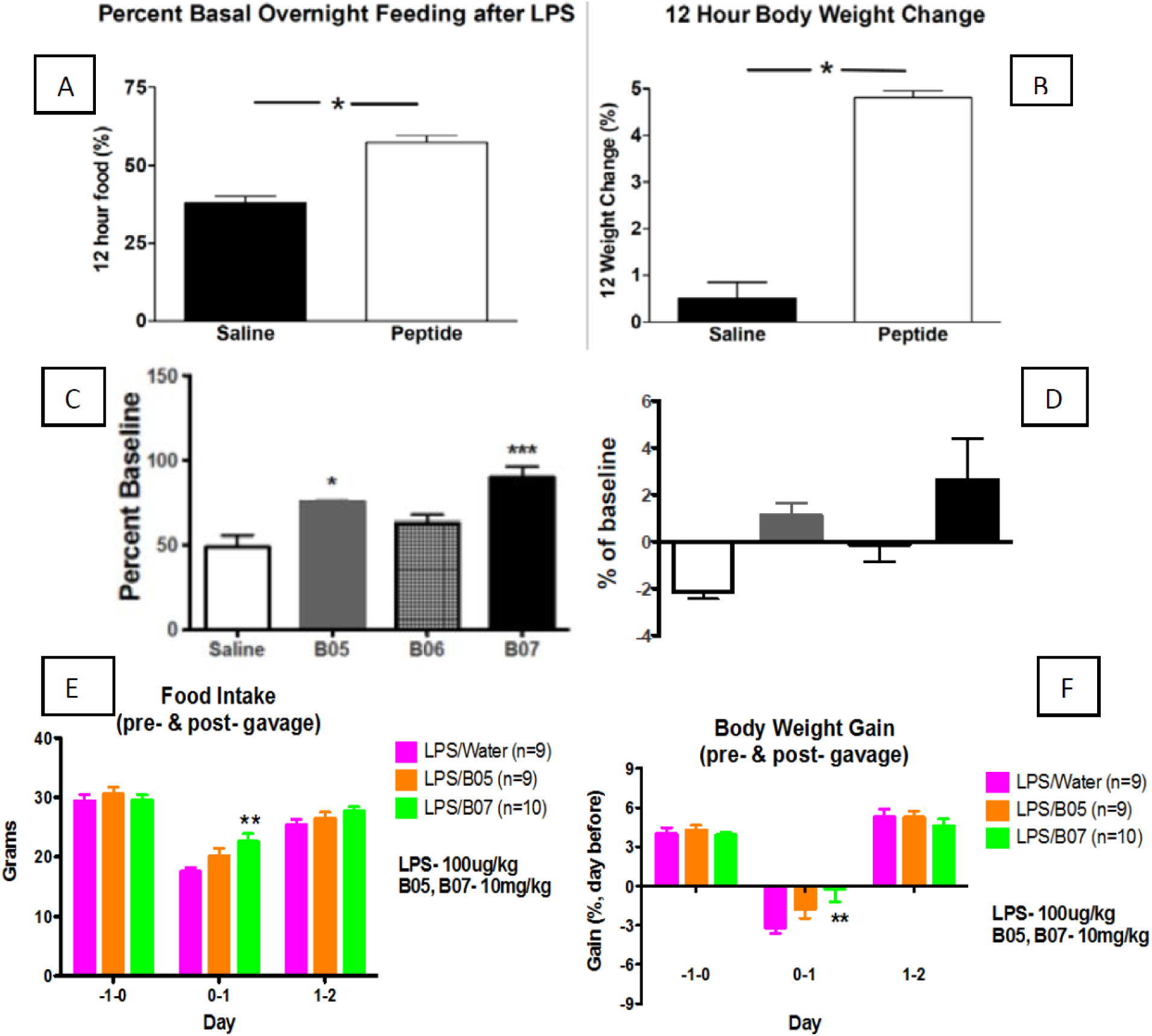
Panels A&B: Shows the effect of parenteral TCMCB02 (peptide) or saline (n=6 for each group) on overnight food consumption and body weight following parenteral LPS administration. Rats eat mainly at night, losing weight during the day and gaining it back at night when they feed. TCMCB02 stimulated feeding (Panel A) compared to a baseline 4 day X 12 hr dark cycle average. Saline controls ate 40% of their baseline, while TCMCB02-treated rats consumed 65%. Saline treated rats gained little weight overnight (Panel B), while TCMCB02-treated rats gained almost 5%. This approached a normal weight gain during their baseline period. Thus, TCMCB02 significantly ameliorated LPS-induced cachexia-anorexia **Panels C&D:** Shows the effect of saline or three different MC4R antagonists in rats given LPS to induce cachexia (n+8/group) on both food intake (Panel C) and body weight (Panel D, as percent of pre-LPS baseline). Measurements were made for 24 hours following the challenge, and drug candidates were given IP at 2mg/kg. Saline controls had anorexia and lost weight. TCMCB05 & 07 stimulated appetite and produced a significant weight gain. TCMCB06 administered rats ate significantly less food than the other drug candidates, with no weight gain. While the weight gain for food eaten with TCMCB05 and 07 appears less than that in Fig 1B, the latter data were recorded at the end of the 12 hr dark cycle when eating occurred. The data in panel D is 24 hour data, which includes the dark/feeding phase and the subsequent 12 hrs of light cycle, when weight loss occurs. * p<0.05, ***p<0.001 **Panels E&F**: Shows the effect of a LPS injection in rats, just before dark cycle, on both food intake and body weight (as percent of pre-LPS baseline). Measurements were made for 24 hours following the challenge, and drug candidates were given IP at 0.2mg/kg. Saline controls had anorexia and lost weight. TCMCB05 & 07 stimulated appetite and produced a significant weight gain. TCMCB06 ate significantly less food than the other drug candidates, with no weight gain. While the weight gain for food eaten in TCMCB05 & 07 appears less than that in Panel B, the latter data were recorded at the end of the 12 dark cycle when eating occurred. These data are 24 hour data, including the the dark/feeding phase and the subsequent 12 hours of light cycle, when weight loss occurs. * p<0.05, ***p<0.001.

***Ac****-Nle-c[Asp-Pro-D-Nal(2′)-Arg-Trp-Lys]-β-Pro-β^2^-Val-****NH*_*2*_** (TCMCB05) and ***Ac***-Nle-c[Asp-Pro-D-Nal(2′)-Arg-Trp-Lys]-β-Pro-β^3^-Val-***NH*_*2*_** (TCMCB06): TCMCB05 and 06 are analogs of TCMCB02, but with metabolism-resistant β amino acid residues (Zubrzak et al., 2007) in their C-terminal dipeptide. The difference between these two molecules is the shifting of either the valine α-carbon amino (TCMCB05) or carboxyl (TCMCB06) groups to the β-carbon. Only TCMCB05 showed parenteral anticachexia activity, compared to TCMCB06 or a saline control (Fig. 1C&D). However, TCMCB05 did not produce significant therapeutic anticachexia activity when given orally (Fig. 1E&F).

***Ac****-Nle-c[Asp-Pro-D-Nal(2′)-Arg-Trp-Lys]-D-Val-D-Pro-****NH*_*2*_** (TCMCB07): We tested the effect of reversing the C-terminal dipeptide sequence of TCMCB02. The resulting peptide, TCMCB07, had parenteral anticachexia activity (Fig 1C&D), and was subsequently tested and shown to have oral anticachexia activity (Fig 1E&F). This peptide was designated as our lead anticachexia drug for further investigation.

#### TCMCB07 Treatment in Rat Cancer Cachexia

We examined the effects of two daily doses of TCMCB07 [total daily treatment of 1.5 mg (low/L; LB07) or 3mg (High/H; HB07) /kg BW/day] in the Lewis sarcoma rat model. We used SC drug administration since i) it is difficult to gavage an untreated-tumor bearing rat due to lethargy, and ii) cachexia adversely affects intestinal absorption (Klein et al., 2013, Puppa et al., 2011).

In Fig 2, the upper panel (A) shows summary drug treatment effects on food intake. Both drug treated groups’ (HB07 and LB07) food intake were not different than Sham/saline controls, and all these groups had significantly greater food intake than the Tumor/saline group.

**Fig 2.**
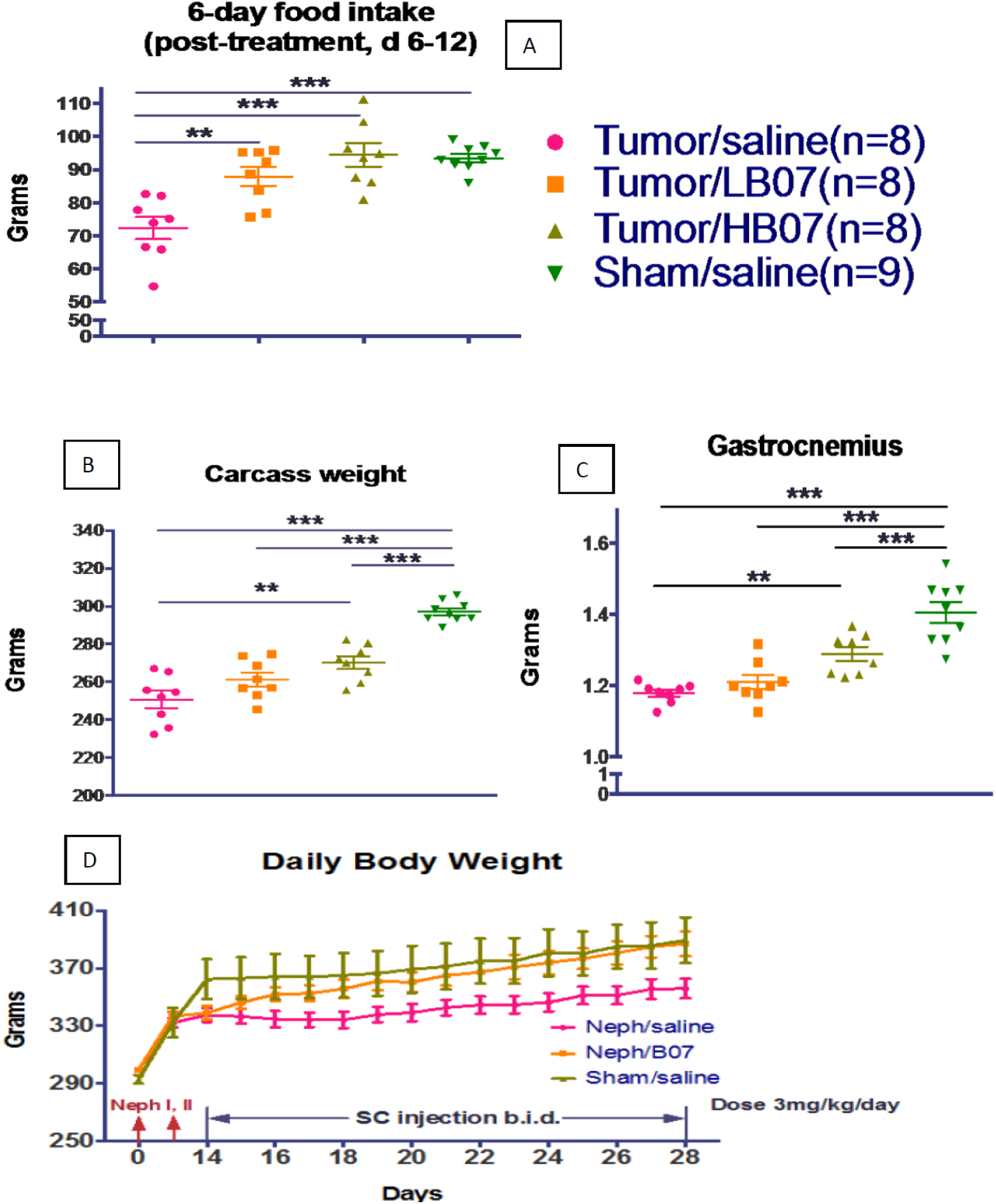
shows the effect of TCMCB07 on two forms of cachexia in rat models. **Upper Panel-A** show summary 6-Day food intake during Lewis Sarcoma cachexia. **Middle Panels-B & C** show terminal carcass weights after the tumor was dissected out, and a skeletal muscle sample, representative of lean body mass. In high and low dose TCMCB07 tumor-bearing groups (HB07 & LB07) there was a drug-dose independent normalization of food intake to control (Sham/Saline) levels. However, the drug-mediated increase in body weight and muscle mass was dose-dependent; with the lower drug dose producing an increase. in weight that was intermediate between the higher drug dose and the untreated (Tumor/saline) group. **Lower Panel-D** shows daily BW during the course of reduced renal mass/uremic cachexia [80% reduction of renal mass (Deboer et al., 2008)] in rats. This results in a chronic renal failure/uremic condition with cachexia. Neph I & II (red arrows on X axis) refer to a unilateral 2/3 reduction in renal mass, followed (7 days later) by removal of the contralateral kidney Neph/Saline BWs (n=11) were compared to the reduced renal mass + TCMCB07 group (Neph/B07; n=11), or a sham operated group (Sham/saline; n=6)). TCMCB07 (or vehicle) administration was begun 1 week after Neph II, when both nephrectomized groups’ BWs were significantly below the sham operated group. Over a 2 week period TCMCB07 treatment (b.i.d. = 2X daily) increased BW to a point where it was equivalent to the Sham/Saline group.

Carcass weights (body weight after the tumor was removed) were significantly depressed in the Tumor/saline group compared to the Sham-saline group (Fig 2B, left middle panel). There were no between group tumor weight differences, indicating that treatment had no effect on tumor growth (data not shown). The HB07 treated group had carcass weights that were significantly greater than the Tumor-Saline group, but were still lower than the Sham-saline group. The LB07 group was not different from either the high dose or the tumor-saline groups; i.e., it was intermediate; even though its summary food intake was not different than the HB07 group (Fig. 2A, upper panel). Gastrocnemius muscle (a lean body mass sample, Fig. 2C, right middle panel) showed the identical pattern as carcass weight, suggesting carcass differences were representative of lean body mass.

#### TCMCB07 Treatment in Rat Reduced Renal Mass Cachexia

We tested the effects of SC TCMCB07 (total dose of 3mg/kg/day) on BW in a reduced renal mass (uremic) cachexia model (Fig 2D, bottom panel). Nephrectomy lowered BWs, with no BW differences between the uremic rats destined for drug treatment versus uremic controls. However, after 2 weeks of drug treatment TCMCB07 significantly increased BW in uremic rats (Neph/B07 group) compared to the vehicle treated uremic rats (Neph/saline group). At that time, the TCMCB07 treated (Neph/B07) group’s BWs were not significantly different than the sham-operated (Sham/saline) group’s BW.

### Pharmacokinetics of TCMCB07 in Rats

The time course of SC administration-induced blood levels of TCMCB07, and its biliary elimination, were evaluated in an anesthetized rat model. Plasma and biliary (cannulated common bile duct) TCMCB07 levels were measured in a group of rats (n=4) following a 3mg/kg SC injection of the peptide. For the first 150 min. both plasma and bile concentrations of peptide rose in a roughly parallel fashion, with the biliary levels displaced by approximately 15-30 minutes (Fig. 3). After 150 minutes, plasma concentrations of the peptide began to fall, while the amount of peptide in bile continued to increase.

**Fig 3:**
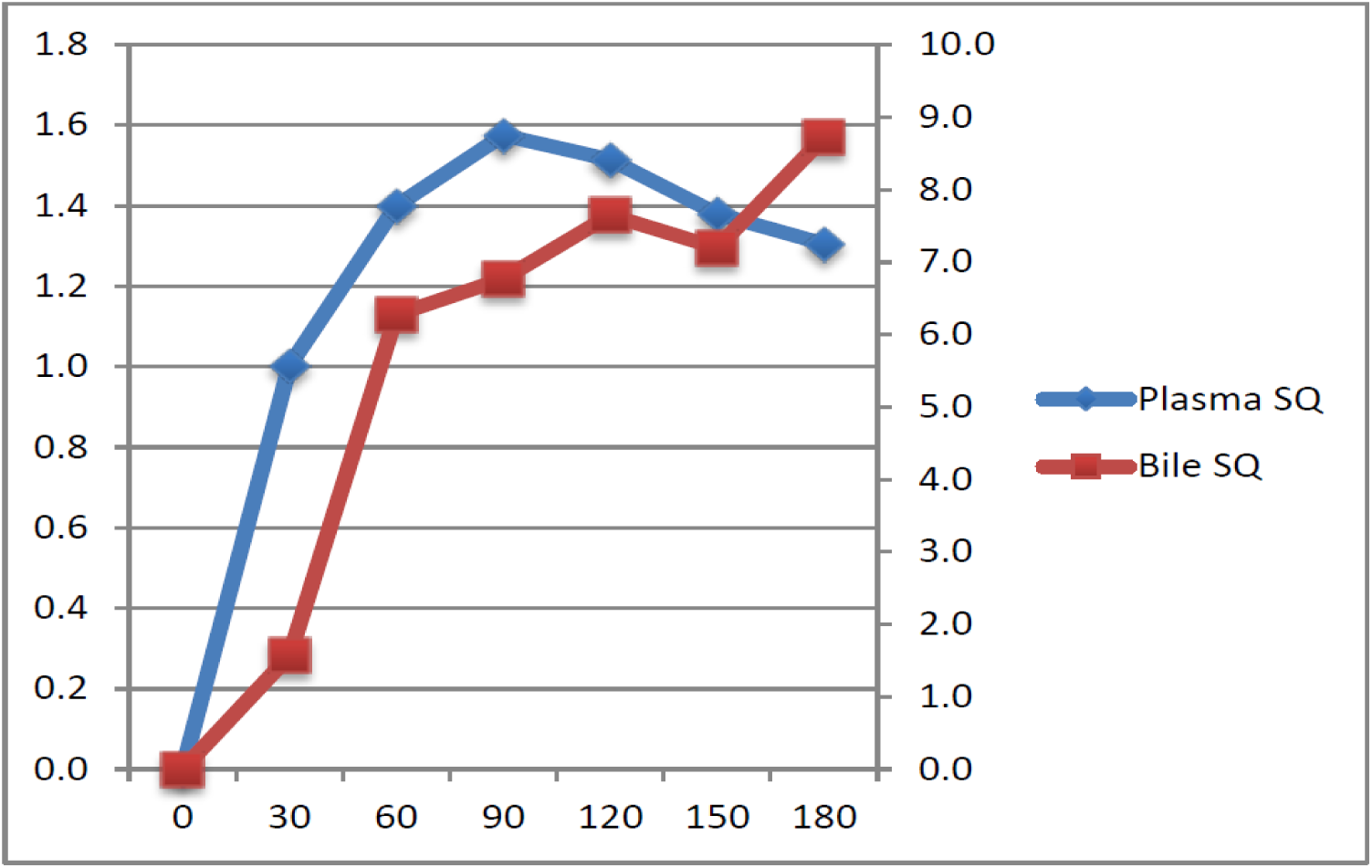
Following a 3mg/kg SC dose of TCMCB07 in anesthetized rats, drug levels were measured via a cannulated femoral vein and cannulated common bile duct. HPLC measured TCMCB07 levels in plasma (left vertical axis in µg/ml) and bile (right vertical axis in µg produced every 10 min). The appearance of the peptide in blood precedes that in bile. The appearance of the peptide in blood precedes its appearance in bile, with the subsequent increases of the peptide in these fluids roughly paralleling each other for the first 90 minutes. The increase of peptide in bile, as plasma levels fall, may reflect a lag in hepatic extraction of peptide from blood vs. peptide secretion into bile.

### Development of an Angiotensin^1-7^ Analog

Angiotensin^1-7^ is an anticancer peptide (Krishnan et al., 2013) whose lack of drug-like properties has inhibited clinical development. Structural examination of angiotensin**^1-7^** (Asp-Arg-Val-Tyr-Ile-His-Pro-OH) revealed a primary sequence that was amenable to vector formation by our platform technology. Thus, we added a Lys-dVal-dPro-amide sequence to the C-terminus, producing a C-terminal D-Val-D-Pro dipeptide and allowing lactam cyclization between Asp**^1^** and Lys**^8^**. The resulting analog, TCANG-05 [Ac-Nle-**c**(Asp-Arg-Val-Tyr-Ile-His-Pro-Lys)-D-Val-D-Pro-NH2], had greatly enhanced stability compared to angiotensin**^1-7^**, using *in vitro* and *in vivo* assays, and was orally active (Gallagher et al., 2016) – see Discussion.

### Development of a Chimeric Vector for an Antiviral Peptide Drug

Gly-Gly-His-Tyr-D-Arg-Phe-Lys was designed as a hybrid of a metal binding domain (GGH) with a hepatitis C virus (HCV) RNA binding peptide. The goal was to produce catalytic destruction of the HCV target (Bradford and Cowan, 2012). While the antiviral activity of this peptide was confirmed in a cellular HCV replicon assay (Bradford and Cowan, 2012), the peptide (Metallodrug 2-Cu) did not have drug-like characteristics (Table 3). We therefore designed analogs of Metallodrug 2-Cu, in which the Gly-Gly-His sequence was linked to a model peptide transport vector whose structure was analogous to the HCV RNA binding peptide.

**Table 3:**
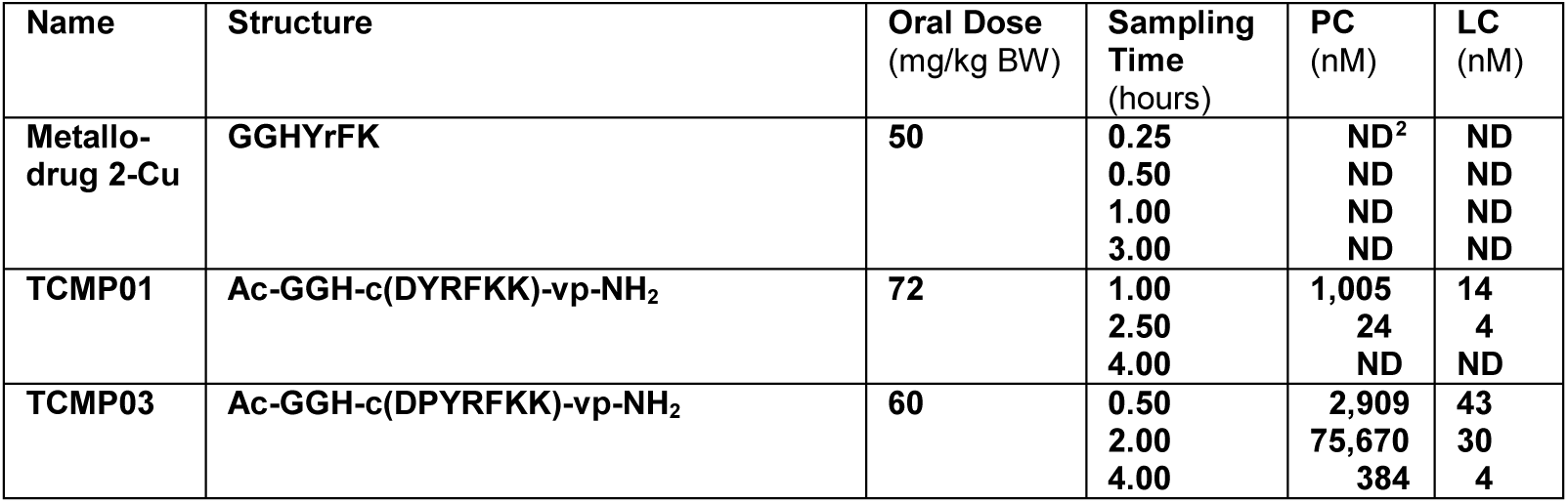
Development of an Orally Active Anti-Hepatitis C Peptide. Shows the sequence of anti-hepatitis C peptides produced and evaluated. Metallodrug 2-Cu was used as the model for analog production. The placement of a prolyl-aromatic residue sequence within the cyclized region of the molecule enhanced oral activity. Sampling Time=number of hours after the administration of each peptide for its measurement in plasma and liver, PC=plasma concentration, LC=liver (anti-HCV target tissue) concentration. Data represent averages of results from duplicate experiments. TCMP01 and 03 are the peptides produced to increasingly conform to the platform technology (see Discussion).

| Name | Structure | Oral Dose<br>(mg/kg BW) | Sampling<br>Time<br>(hours) | PC<br>(nM) | LC<br>(nM) |
| --- | --- | --- | --- | --- | --- |
| Metallo-<br>drug 2-Cu | GGHYrFK | 50 | 0.25 | ND<sup>2</sup> | ND |
|  |  |  | 0.50 | ND | ND |
|  |  |  | 1.00 | ND | ND |
|  |  |  | 3.00 | ND | ND |
| TCMP01 | Ac-GGH-c(DYRFKK)-vp-NH<sub>2</sub> | 72 | 1.00 | 1,005 | 14 |
|  |  |  | 2.50 | 24 | 4 |
|  |  |  | 4.00 | ND | ND |
| TCMP03 | Ac-GGH-c(DPYRFKK)-vp-NH<sub>2</sub> | 60 | 0.50 | 2,909 | 43 |
|  |  |  | 2.00 | 75,670 | 30 |
|  |  |  | 4.00 | 384 | 4 |

We began with a Tyr-Arg-Phe-Lys sequence, analogous to the HCV RNA binding domain. We then added flanking Asp and Lys residues for lactam cyclization of the peptide, and a C-terminal D-Val-D-Pro motif. We synthesized a peptide containing this transport vector with the Gly-Gly-His metal binding sequence at its amino terminus (TCMP01, Table 3). TCMP01 maintained significant anti-hepatitis C activity, as reflected in standard solution assays (Bradford and Cowan, 2012, Bradford et al., 2014) as well as improved drug-like properties; including extended plasma half-life, and hepatic uptake compared to Metallodrug 2-Cu (Table 3). However, while the oral bioavailability of TCMP01 was an improvement on its linear analog, it was still relatively limited. Further improvements in circulating half-life, oral activity, and hepatic uptake were produced following the addition of a Pro residue on the amino side of the Tyr residue (a Pro-aromatic motif; see Table 1) in the transport vector (TCMP03, Table 3).

Similar to TCMP01, TCMP03 displayed equivalent solution anti-hepatitis C activity. Relative to the parent peptide, Metallo-drug 2-Cu, both TCMP01 and 03 had solution antihepatitis activity within a factor of two. However, the latter peptides showed a 14-43-fold enhancement of hepatic uptake (Table 3, LC column).

### NMR Spectroscopy of TCMCB07

We investigated the structural characteristics of our lead MC anticachexia agent, TCMCB07, using solution NMR spectroscopy and compared it to a close analog TCMCB03. The latter peptide has a His for Pro substitution at position 3 of TCMCB07, and lacks parenteral anticachexia activity (i.e., does not cross the BBB; data not shown).

NMR spectra of TCMCB07 provided NOE-based distances, J-coupling information on dihedral angles, and temperature coefficients consistent with backbone amide groups in hydrogen bonds.

The NOE-based close contacts and dihedral information were used to restrain molecular dynamics (MD) simulations. The NMR-restrained structural ensemble is evidence that the cyclized region of TCMCB07 forms a β hairpin with the Pro**^3^** and D-Nal**^4^** being central in the β-turn (Fig. 4). Though NOEs support a β-turn in TCMCB03 with the His^3^ substitution (Supplemental Information), **^15^**N NMR (T2) relaxation data suggest the backbone of TCMCB03 to be more flexible than that of TCMCB07 (see Supplemental Information). Thus, the His**^3^** substitution destablilizes the β hairpin structure.

**Fig 4:**
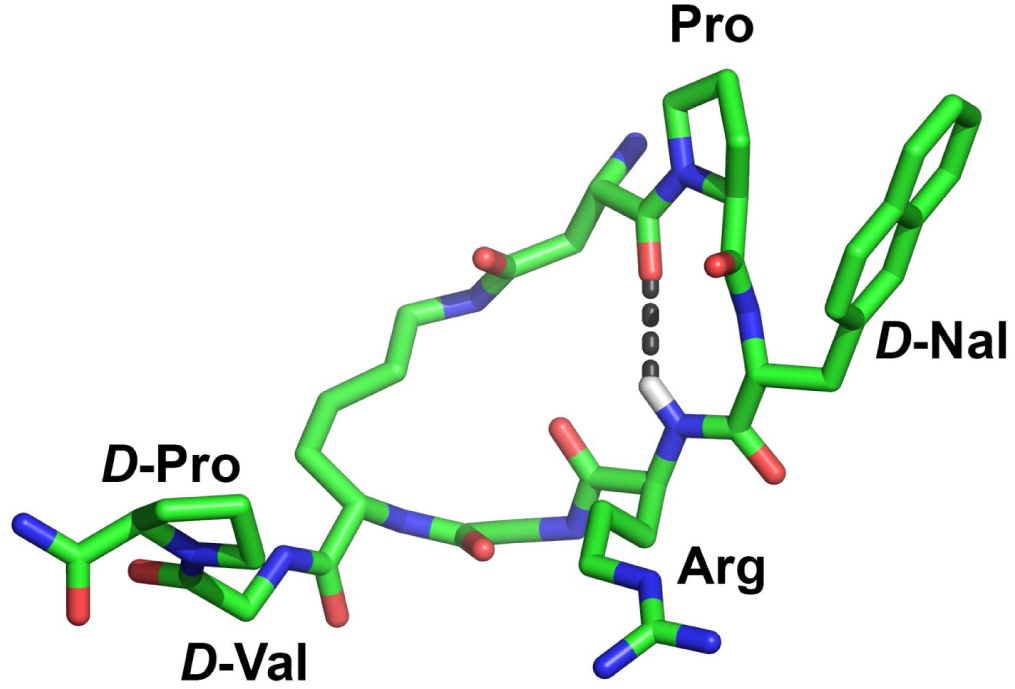
Structural model of TCMCB07 simulated by molecular dynamics restrained by NMR measurements. The estimates of close contacts are based on NOEs and the torsion angle ranges upon J-couplings. This conformation agrees well with the NOEs and J-couplings, according to lowest violations of the restraints. The dashed line marks the probable hydrogen bond between the Arg^5^ (N-and Asp^2^ (C=O). This closes the β hairpin turn centered at Pro^3^-Nal^4^ in the cyclized region of the molecule. See Supplemental Data Items, Tables 1S-5S and Fig. 5S.

The His**^3^**-induced changes in NMR peak positions are greatest between Arg**^5^** through Val**^8^**, consistent with the most disruption of the peptide structure occurring in this segment. The NMR spectra of covalently homogeneous TCMCB03 also possess two sets of NMR peaks in both aqueous solution and DMSO-d6 solution; evidence that TCMCB03 exists in two sets of conformers with similar populations (approximately 1:1 ratio). The equilibrium between the conformers does not shift within the range of temperature and pH changes tested. Therefore the more rigid β hairpin secondary structure of TCMCB07 may correlate with its bioactivity.

## Discussion

Vector mediated drug delivery has been used to transport peptides and other molecules across the BBB (Bockenhoff et al., 2014). Compounds (“vectors”) that undergo BBB transport by active or transcytotic mechanisms are covalently linked to potentially therapeutic agents that lack this ability. This produces chimeric molecules with the BBB transport characteristics of the vector, while expressing therapeutic activity. We developed a vector technology that goes beyond simply BBB transport, producing peptides with oral activity, extended *in vivo* half-life, as well as BBB transport.

We chose the central MC system in cachexia as our initial therapeutic target for developing a drug-like peptide. MC4R antagonists or genetic disruption of the MC4R has consistently enhanced weight gain in all vertebrates tested; zebrafish (Song and Cone, 2007), through humans (Vaisse et al., 1998). I*n vivo* structure:activity studies were modeled around an established anticachexia MC4R antagonist peptide that had apparent BBB transport activity; PG932 (Table 1). We noted that PG932’s structure included two dipeptide features that differentiated it from an analogous MC4R antagonist peptide (SHU9119) that lacked BBB transport (Sutton et al., 2008). These included a proline (adjacent to an aromatic residue) within the cyclized region of the sequence, and a C-terminus composed of non-polar residues from the flanking region of α-MSH; Pro-Val (Eberle and Schwyzer, 1976). Similar sequence motifs are present in non-MC peptides that express *in vivo* transport (Table 1; peptides #2, 3, 4, and 5-12). We incorporated metabolism-resistant “D” enantiomers and β analogs of the residues in the C-terminal dipeptide of PG932, in an attempt to produce an analog with oral anticachexia activity by enhancing *in vivo* half-life.

Several MC4R peptide antagonists were screened for anticachexia activity to a bacterial toxin/LPS challenge using parenteral and oral routes of peptide administration. This approach initially identified 3 peptides with parenteral anticachexia activity (i.e., able to cross the BBB). While the D-Pro-D-Val derivative of PG932 was effective via the parenteral route (Fig. 1 A&B), there was no evidence for oral activity.

Substituting different forms of β amino acid residues in the C-terminal dipeptide demonstrated the highly specific structural requirements of the C-terminal dipeptide for transport. Only the MC analog with the β**^2^** form of valine in the C-terminal dipeptide (TCMCB05) maintained parenteral/BBB transport activity (Figure 1 C&D). TCMCB06, with a C-terminal β**^3^** valine substitution, had no parenteral activity (Figure 1 C&D): i.e., it did not cross the BBB in therapeutically significant quantities. Structurally related cyclic peptides with highly disparate trans-epithelial permeability are suggestive of carrier-mediated transport (Marelli et al., 2015).

The C-terminal dipeptide sequence of peptide #2 in Table 1 contains a valine analog residue [Tle=tert(iary) leucine=3-methyl valine] in the penultimate position. This suggested the testing of a reversal of the C-terminal dipeptide in TCMCB02 (D-Pro-D-Val) to D-Val-D-Pro (TCMCB07). This dipeptide sequence placed a D-Pro residue on the carboxyl side of a nonpolar residue, analogous to the structures of peptides #6-12 in Table 1. Reversing the TCMCB02 C-terminal dipeptide sequence also tested the specificity of this C-terminal motif for transport. TCMCB07, had oral anti-cachexia activity (implying the maintenance of BBB transport; Fig 1E-F, as well as gastrointestinal transport). Other C-terminal dipeptide residues may also produce some form of carrier transport, including leucine and tertiary leucine (Table 1, peptide #2), as well as phenylalanine and tryptophan (Table 1, peptides #2-4 and 6-12).

Based on these data, the critical requirements for producing a drug-like MC peptide were hypothesized to be a proline residue in close proximity to an aromatic residue (usually within a cyclized region of the peptide) and a dipeptide sequence (at the C-terminus or within a head-to-tail cyclized structure) composed of non-polar residues; preferably D-Val-D-Pro. We investigated the general applicability of these structural modifications for their ability to produce drug-like activity by applying them to non-MC peptides with potential therapeutic activities, but lacking the drug-like properties required for clinical utility.

Angiotensin**^1-7^** is an anticancer/MAS receptor peptide agonist (Krishnan et al., 2013, Menon et al., 2007, Soto-Pantoja et al., 2009, Gallagher et al., 2011). Like many peptides, angiotensin**^1-7^** shows poor PK properties, with a plasma half-life of ∼30 minutes (Iusuf et al., 2008). This has been a major impediment to the clinical development of this peptide, despite the evidence for its efficacy in human cancers (Petty et al., 2009, Rodgers et al., 2006). Using TCMCB07 as a model for the development of a drug-like peptide, we designed and synthesized a cyclized analog of angiotensin**^1-7^:** TCANG05. The resulting peptide had the two previously defined essential features for drug-like properties: a cyclized region containing a Pro adjacent to an aromatic (His) residue, and a D-Val-D-Pro C-terminal dipeptide.

Gallagher et al. assessed the effects of angiotensin**^1-7^** and TCANG05 on cell growth in cultures of human A549 lung and MDA-MB-231 breast cancer cells over 6 days (Gallagher et al., 2016). Significant growth inhibition by angiotensin**^1-7^** required daily dosing of the cultures. In contrast, a single dose of TCANG05 (on day 1) produced growth inhibition for 6 days, suggesting enhanced peptide stability. In a separate experiment, angiotensin**^1-7^** had a 30 min plasma half-life, while the half-life of TCANG05 was 50 hours.

The *in vivo* activity of TCANG05 was tested by oral gavage in an athymic mouse model bearing an orthotopic human breast cancer cell line transplant (cell line MDA-MB-231). Angiotensin**^1-7^** was not compared in this study since it is not orally active. TCANG05 caused a significant reduction in tumor volume and tumor weight; e.g. a 60 µg/kg/day dose reduced tumor volume 57% compared to saline gavaged mice (Gallagher et al., 2016). Further, TCANG05 did not affect mouse BW, or the size of its heart or kidneys.

In addition to using modifications in a peptide’s structure to produce a transport vector, another vector-based solution is to link a potentially therapeutic peptide to a transport vector to produce a hybrid or chimeric peptide. For example, we were presented with a hybrid peptide construct (Table 3, Metallodrug 2-Cu) consisting of a GGH motif with a hepatitis C RNA binding/cell penetrating motif (YrFK) at its amino terminus. The GGH sequence was designed to carry metal ions to a viral target, producing catalytic irreversible damage (Bradford and Cowan, 2012). A hybrid containing the GGH and YrFK sequences could potentially bind to and destroy the hepatitis C virus. However, the hybrid peptide was lacking in several critical drug-like properties; no oral availability, a short circulating half-life, and poor target tissue (liver) uptake. While the metal ion binding activity of the GGH sequence would not withstand cyclization, examination of Metallodrug 2-Cu’s structure suggested that the RNA binding motif was potentially amenable to derivatization by our technology.

Our first analog placed an Asp residue between Gly**^3^** and His**^4^**, replaced the D-Arg**^5^** with an L-Arg, and added a Lys-D-Val-D-Pro-amide to the C-terminus. The peptide was then lactam cyclized between Asp**^4^** and Lys**^8^.** This peptide (TCMP01, Table 3) had some degree of oral activity, but not to the extent to where it was a potentially therapeutic agent *in vivo*. We then synthesized a second derivative in which a Pro residue was placed between Asp**^4^** and Tyr**^5^**, producing a Pro-aromatic residue motif within the cyclized region. This peptide (TCMP03) had 46-fold more oral activity than TCMP01, with a 4-fold enhancement of hepatic (target tissue) uptake (Table 3). These analogs confirm the importance of the C-terminal D-Val-D-Pro, and a Pro-aromatic dipeptide for producing drug-like properties.

Thus, applying the structural requirements for producing a drug-like MC peptide to unrelated peptides produced similar *in vivo* properties. These data provide evidence that for the utility of these derivatizations in the production of drug-like peptides.

Previous reports suggested that transport of cyclized peptides is enhanced when derivatized to a stable secondary structure (Beck et al., 2012, Hess et al., 2007). However, attempts to use this concept in a predictive fashion to produce drug-like properties in peptides failed (Hess et al., 2007).To more fully define the structural basis for the transport properties of our peptide analogs, we used TCMCB07 as a model peptide with transport activity, and performed solution NMR structural studies. As a negative control for these experiments, we used an analog of TCMCB07 in which Pro**^3^** was replaced by a His residue (TCMCB03). The MC antagonist pharmacophore in TCMCB03 (His-D-Nal-Arg-Trp) is identical to that in SHU9119, an established MC4R antagonist. However, SHU9119 has no transport activity, requiring intracerebroventricular administration for use as an anticachexia agent (Sutton et al., 2008). TCMCB03 has a similar lack of parenteral anticachexia activity (i.e. no BBB transport).

The cyclized region of TCMCB07 forms a β hairpin, centered at the Pro-Nal sequence (Fig. 4). This is consistent with the evidence that aromatic and proline residues contribute to peptide stabilization into a β hairpin (Meyer et al., 2013, Haque and Gellman, 1997). The NMR-based structural model (Fig. 4) suggests a hydrogen bond between Asp**^2^** (C=O) and Arg**^5^** (N-H) that contributes to ring stabilization. In contrast, NMR analysis of TCMCB03, with a His for Pro**^3^** substitution, revealed a more dynamic peptide (Fig. 5S), with two conformational populations. This is consistent with the evidence for multiple conformations in solution for SHU9119 (Victoria Silva Elipe et al., 2003).Thus, the His for Pro substitution may have a global disruptive impact on the secondary structure of this peptide. This lack of structural stability could be related to the absence of transport activity in TCMCB03.

Review of the literature revealed a candidate carrier system that could account for the transport actions associated with TCMCB07. Cyclosomatostatin and structurally related peptides (Table 1; peptides # 6-12) undergo blood-to-bile active transport (Ziegler et al., 1985, Petzinger et al., 1983, Frimmer and Ziegler, 1988). The mechanism for this appears to be direct competition at a hepatocyte anionic active transporter for bile salts (Ziegler et al., 1991).

Most of the peptides in Table 1 possess at least one of our requirements for drug-like transport activity; 10 of these 12 had a Pro-aromatic dipeptide within a cyclized or internal region of the molecule. Of the 10 peptides, all had at least one additional dipeptide motif composed of non-polar residues including valine or valine analogs, and in 3 of the 10 the non-polar dipeptide was a Val-Pro or Pro-Val sequence. Another analogy is that one of the *in vivo* mechanisms for clearence of TCMCB07 is hepatic blood-to-bile transport (Fig. 3).

More recent work suggests that both the OATP and OAT families of transporters have members with bile acid/cyclic peptide transport capabilities. These transporters are located in the intestine and BBB (blood to CSF transport direction), and are therefore potential candidates for the transport system used by peptides produced by our platform technology (Sekine et al., 1998, Dalvi, 2014).

In summary, our technology provides at least two approaches for imparting drug-like properties to a peptide of interest: (i) converting a peptide into a transport vector, or (ii) covalent linkage of a therapeutic peptide to a model peptide vector produced by our platform. It should be noted that the latter approach suggests the possibility of designing a peptide drug that expresses multiple therapeutic activities. As proof of this concept, we produced a chimeric peptide consisting of a cyclic cationic-aromatic anti-microbial peptide [e.g. (Dathe et al., 2004)] and a linear anti-biofilm peptide [e.g. (Dean et al., 2011)]. While antimicrobial activity of peptides is enhanced by cyclization, anti-biofilm peptides cannot be cyclized and maintain activity, since they require a helical secondary structure (Dean et al., 2011). This hybrid peptide expressed both activities (unpublished data). The generalizable nature of our platform awaits further testing in additional classes of peptides with therapeutic activities.

## EXPERIMENTAL PROCEDURES

Non-commercially available peptide sequences were designed by Tensive Controls’ scientists, and synthesized by solid phase technique at the University of Missouri Peptide Synthesis Core, or under cGMP conditions by CPC, Inc. (Sunnyvale, CA). Custom made peptides can be expensive to produce, precluding their general availability to other scientist. However, Tensive Controls, Inc. will agree to a limited license, to allow the synthesis of milligram quantities of any peptide disclosed in this publication, for purposes of confirmation and other non-commercial uses.

### Routes of Administration

Peptides or vehicle (0.9% saline) were given by: subcutaneous (SC) administration in 0.45% saline or dissolved in water in an oral gavage.

### Therapeutic Assays/Models

All animal protocols were approved by the Institutional Animal Care and Use Committees of the University of Missouri-Columbia, Oregon Health Sciences University, and Ohio State University. Protocols were in compliance with the National Institutes of Health and U.S. Department of Agriculture Guidelines for Care and Use of Animals in Research.

i. *Normal Rat and Mouse Studies*
  a. *TCMCB07 Pharmacokinetic Studies:* Studies were performed in isoflurane-anesthetized Sprague-Dawley rats with femoral vein catheters, and (in some experiments) a common bile duct catheter (see Supplemental Experimental Procedures for further details).
  b. *Antiviral Peptide Studies: P*eptides were given by oral gavage to female mice. (see Supplemental Experimental Procedures for further details).
ii. *Rat Cachexia Models:* Note: “cachexia” (wasting of body mass) is technically termed “cachexia-anorexia syndrome”, since loss of appetite is usually associated with the condition. However, loss of body mass is the most relevant metric in assessing cachexia, since it reflects the hypermetabolic state that is associated with cachexia-induced multiorgan pathology (Fukuda et al., 2009, Klein et al., 2013, Amitani et al.,2013). Thus, unless specifically noted, we use the term anti-cachexia to refer to the preservation of body weight (BW).
  a. *Endotoxin:* Lipopolysaccharide (LPS) was used to produce endotoxin-induced cachexia. (See Supplemental Experimental Procedures for further details).
  b. *Cancer*: The Lewis sarcoma rat model is a methyl-cholanthrene-induced sarcoma that produces significant cachexia about 6-8 days after tumor implantation (Popp et al., 1982). The tumor can be removed (it does not metastasize), to compare true BW changes without the confounding tumor mass (Popp et al., 1982). (See Supplemental Experimental Procedures for further details).
  c. *Reduced Renal Mass*: A reduced renal mass model was produced as previously described (Deboer et al., 2008) (see Supplemental Experimental Procedures for further details).

### Analytical Procedures

i. *Analytical Method for Melanocortins in Rats:* We developed an assay for MC peptide antagonists in body fluids using acetonitrile extraction and reverse-phase HPLC separation (see Supplemental Experimental Procedures for further details).
ii. *Analytical Method for Antiviral Peptides in Mice*: Chromatographic and mass spectrometric experiments were conducted on an Agilent 1260 Infinity UHPLC system, and Agilent 6460 Triple Quadrupole LC/MS with Jetstream technology and electrospray source, respectively (see Supplemental Experimental Procedures for further details).

### NMR & Molecular Modeling Methods

i. All NMR experiments were carried out on an 800MHz Bruker Avance III spectrometer with TCI cryoprobe at 300K. 1D proton spectra were acquired between 280 and 320K to measure the temperature coefficients of the amide protons. 2D NMR DQF-COSY, TOCSY and NOESY spectra were recorded in all solvents for assignment (see Supplemental Experimental Procedures for further details).
ii. *Molecular Modeling:* The computational protocol for structural determination in solution consisted of restrained molecular dynamics (RMD) simulations, energy minimization and cluster analysis (see Supplemental Experimental Procedures for further details).

## Technical Assistance

We acknowledge the scientific contributions of Dr. Daniel Marks and his laboratory staff in the assay of anticachexia activity.

## Author Contributions

### Participated in research design

Gruber, Callahan, Cowan, Jiang, and Van Doren *Conducted experiments:* Gruber, Callahan, Gallazzi, Jiang, Cowan, Qi, Pearson, Ross, and Wachnowsky

### Contributed new reagents or analytical tools

Gruber, and Cowan

### Performed data analysis

Callahan, Cowan, Gallazzi, Jiang, and Cowan

### Wrote or contributed to writing of manuscript

Gruber, Callahan, Cowan, Jiang, Van Doren

## Footnotes

1 Subscript “c” before parentheses indicates cyclized sequence within

2 ND: Not Detectable

